# Assessing the effect of experimental evolution under combined thermal-nutritional stress on larval thermotolerance and thermal plasticity in *Drosophila melanogaster*

**DOI:** 10.1101/2024.06.12.598748

**Authors:** Yeuk Man Movis Choy, Teresa Kutz, Fiona E. Cockerell, Emily Lombardi, Sandra Hangartner, Christen Mirth, Carla M. Sgrò

## Abstract

Animals commonly face combinations of thermal and nutritional stress in nature, which will intensify under climate change. While genetic adaptation is necessary to buffer long-term stress, it’s unclear whether adaptation to combined stress can occur without compromising viability and thermal plasticity. We tested larval thermotolerance and thermal plasticity in *Drosophila melanogaster* selected under different temperatures (18°C, 25°C, and 28°C) and diets (standard, diluted, and low-protein:high-carbohydrate [P:C]). Basal larval cold tolerance was affected by both protein concentration and temperature; larvae evolved higher basal cold tolerance on the diluted and low P:C diets at 18°C and 28°C. Hardening increased cold tolerance for most lines, except those selected at 18°C and 28°C on low P:C diets and at 25°C on standard diets. Basal larval heat tolerance was affected by selection temperature; selection at 25°C increased heat tolerance. An interaction between selection temperature, selection diet, and hardening treatment affected larval heat tolerance; hardening reduced heat tolerance in most lines, except those selected at 25°C on low P:C diets and at 28°C on standard diets. Our results suggest that adaptation to combined stress allows basal cold tolerance and its plasticity to co-evolve, but not heat tolerance, highlighting ectotherm’s vulnerability to long-term climate change.

## Introduction

Anthropogenic climate change is increasing the frequency of climatic extremes (Thomas *et al*. 2004; Diffenbaugh and Field 2013; Wiens 2016). Combined shifts in atmospheric CO2, precipitation, and temperature (Van Der Putten *et al*. 2010; Pecl *et al*. 2017; Masson-Delmotte *et al*. 2018) will alter plant growth, nutritional quality (Kreuzwieser and Gessler 2010; Lukac *et al*. 2010; Jin *et al*. 2019), and abundance (Van Der Putten *et al*. 2010; Pecl *et al*. 2017). These changes will likely impose increasing thermal and nutritional stress on organisms, both directly and indirectly through changes in microbial communities associated with plant decomposition (Bowes 1993; Richardson *et al*. 2002; Dwyer *et al*. 2007; DaMatta *et al*. 2010; Rosenblatt and Schmitz 2016). Ectothermic herbivores and frugivores, which rely on ambient temperature to maintain physiological performance and function (Hawlena and Schmitz 2010; Walters *et al*. 2012), will be particularly vulnerable to increased thermal and nutritional stress.

Organisms primarily respond to environmental stress by two means, phenotypic plasticity and genetic adaptation (Stillwell *et al*. 2007; Urban *et al*. 2014). Phenotypic plasticity, in which a single genotype produces different phenotypes depending on the environment, is often the initial response to environmental change (Huey *et al*. 2002; Urban *et al*. 2014; Seebacher *et al*. 2015; Sgrò *et al*. 2016). Plasticity allows for immediate (within-generation) responses to environmental change (Padilla and Savedo 2013; Sgrò *et al*. 2016) by shifting behaviours, physiologies, and morphologies that may contribute to population persistence (Chevin *et al*. 2010; Frankino *et al*. 2019; Noble *et al*. 2019), although it may not always be adaptive (Ghalambor *et al*. 2007; Leonard and Lancaster 2020). Nonetheless, an increasingly large body of work is focused on understanding plastic responses to a range of stressors, particularly temperature and nutrition, to better predict population persistence under climate change. Many of these studies have focussed on plastic shifts in thermotolerance (van Heerwaarden *et al*. 2016; van Heerwaarden and Kellermann 2020; Rodrigues and Beldade 2020) since the ability to withstand thermal extremes is a significant predictor of climate sensitivity (Hoffmann *et al*. 2013; Noh *et al*. 2017; Kellermann and van Heerwaarden 2019; Ma *et al*. 2021).

Plasticity in response to temperature and nutrition individually has been widely reported, particularly in insects (Lee, 1989; Andersen *et al*. 2010; Sgrò *et al*. 2016; Mitchell *et al*. 2017; Ma *et al*. 2021). For example, diets with higher protein increased adult heat tolerance in insects (Andersen *et al*. 2010), while higher dietary carbohydrates facilitate the synthesis of cold-resistant compounds, which in turn enhance cold tolerance (Hazel 1995; Kostal and Simek 1998; Andersen *et al*. 2010; Colinet *et al*. 2013).

Thermal plasticity can arise through acclimation (Rako and Hoffmann 2006; Andersen *et al*. 2010; Kingsolver *et al*. 2016), which involves long-term exposure of days or weeks to temperatures above or below the optimal temperatures but still within the normal viable range of an organism. Acclimation to higher developmental temperatures often increases basal heat tolerance (Kingsolver *et al*. 2016; Kellermann *et al*. 2017; Pumhan *et al*. 2020; Jerbi-Elayed *et al*. 2021; Willot *et al*. 2021), while acclimation to lower developmental temperatures can increase basal cold tolerance (Rako and Hoffmann 2006; Scharf *et al*. 2015; Noh *et al*. 2017).

Hardening, the short-term exposure of minutes or hours to sublethal temperatures, can also induce plastic shifts in thermotolerance (Hoffmann *et al*. 2003; Bowler 2005; Loeschcke and Sørensen 2005; Teets *et al*. 2020; Ma *et al*. 2021). In insects, high- and low-temperature hardening treatments can improve subsequent heat (Chen *et al*. 1991; Sejerkilde *et al*. 2003; Bubliy *et al*. 2012; Manenti *et al*. 2015; Kalra *et al*. 2017) and cold tolerance (Czajka and Lee 1990; Chen *et al*. 1991; Kelty and Lee 2001; Sejerkilde *et al*. 2003; Jensen *et al*. 2007; Jakobs *et al*. 2015; Teets *et al*. 2020), respectively.

Plastic shifts in response to combinations of thermal and nutritional stress have been documented, but to date, such studies have primarily focussed on life history traits rather than thermal tolerance (Polak *et al*. 2004; Bubliy *et al*. 2012; Kutz *et al*. 2019; Chakraborty *et al*. 2020). Interestingly, these studies suggest that plastic responses to combinations of thermal and nutritional stress may actually be non-adaptive. For instance, larvae of *Tenebrio molitor* and *Drosophila melanogaster* developed at higher temperatures in combination with suboptimal diets, had decreased survival (Rho and Lee 2017; Kutz *et al*. 2019).

Despite widespread evidence of thermal plasticity, studies suggest that plasticity is unlikely to fully mitigate the negative consequences of ongoing temperature increases. For example, plasticity was found to have a limited effect on increasing the upper thermal limit of *D. melanogaster* (van Heerwaarden *et al*. 2016). There is also some evidence suggesting that insects with higher basal thermotolerance may be less plastic (Nyamukondiwa *et al*. 2011; Gerken *et al*. 2015; van Heerwaarden *et al*. 2016; Kellermann *et al*. 2017; Noh *et al*. 2017; Kellermann and Sgrò 2018), implying a potential trade-off between plasticity and basal thermotolerance (but see van Heerwaarden *et al*. 2024). Thus, in the context of ongoing climate change, genetic adaptation, while occurring over longer timescales, will be key to the long-term persistence of populations (Potvin and Tousignant 1996; Gienapp *et al*. 2008; Lande 2009).

Reliable predictions concerning whether and how populations might respond to long-term selection imposed by climate change require answers to key questions (Cavicchi *et al*. 1995): Do populations harbour the genetic variation necessary for rapid evolutionary responses to combinations of stressors? Does adaptation to specific thermal and nutritional conditions involve genetic changes in many traits? To what extent does adaptation affect the ability of populations to mount plastic responses to conditions other than those selected?

Experimental studies of evolution in the laboratory (Hoffmann and Harshman 1999; Rose *et al*. 2005) have addressed these questions with respect to genetic adaptation to thermal (Huey *et al*. 1991; Cavicchi *et al*. 1995; Bubliy and Loeschcke 2005) and nutritional (Kolss *et al*. 2009; Kristensen *et al*. 2011; Leftwich *et al*. 2017) stress individually. Cavicchi *et al*. (1995) and Huey *et al*. (1991) selected adult *D. melanogaster* at different non-extreme temperatures and observed rapid evolution in their thermotolerance. Lines adapted to higher temperatures evolved higher heat tolerance (Huey *et al*. 1991; Cavicchi *et al*. 1995), while those adapted to cooler temperatures had faster development at lower temperatures, but their cold tolerance was not assessed (Huey *et al*. 1991). Interestingly, Cavicchi *et al*. (1995) found that thermal adaptation also resulted in shifts in thermal plasticity. Lines evolved at 18°C showed a positive acclimation response (heat tolerance increased) to development at 25°C and 28°C, while 25°C evolved lines showed a positive acclimation response to 25°C, but a negative acclimation response (heat tolerance decreased) to 28°C. Further, the evolved lines also showed changes in their hardening responses, with the 25°C evolved lines having a higher hardening threshold (the temperature that induces a hardening response) compared to the 18°C evolved lines. Notably, the 28°C evolved lines lost the ability to mount a hardening response, and exposure to the highest hardening temperature reduced thermal tolerance. These results suggest that adapting to elevated temperatures may come at the cost of reduced plasticity. However, the effect of thermal adaptation on larval thermotolerance traits was not examined.

Experimental evolution has also been used to assess adaptation to changes in diet. Kolss *et al*. (2009) examined adaptation to nutritional stress in larvae of *D. melanogaster* by exposing them to poor (25% diluted) or control diets throughout development; adults were placed on the control diet every generation. Evolved lines were tested on both poor and control diets to assess whether adaptation was associated with any trade-offs in life-history traits and whether nutritional plasticity had evolved. When tested on the poor diet, the selected lines had higher larval viability than the control lines, but viability remained the same for both lines when tested on the control diet. However, adults of lines selected on poor diets were smaller, less fecund, and less resistant to starvation than the controls. Interestingly, there was no effect of selection or test diet on larval heat or cold tolerance. Similarly, Kristensen *et al*. (2011) used experimental evolution to expose *D. melanogaster* larvae to standard or protein-enriched diets. They found that adaptation to the high-protein diet resulted in reduced larval-adult survival, suggesting that adaptation to changes in nutrition may be limited by trade-offs between life-history traits. However, they did not examine larval or adult thermotolerance. Finally, Leftwich *et al*. (2017) imposed nutritional selection on larvae of the Mediterranean fruitfly and found that lines selected on the low-calorie diet evolved higher egg-adult survival compared to those on the standard diet after 30 generations of selection, when tested on both diets. Once again, larval thermotolerance was not assessed.

Only one study has examined adaptation to both nutritional stress and temperature. Bochdanovits and De Jong (2003) exposed larvae to two temperatures (17.5°C and 27.5°C) and poor (50% diluted) and control diets. Overall, adaptation to the higher temperature resulted in reduced larval survival on both diets. The 17.5°C selected lines evolved larger body size when also adapted to a poor diet, while the 27.5°C evolved lines were smaller on both diets. This suggests that adaptation to combinations of temperature and nutrition can result in trade-offs between adult size and larval survival, and potentially other traits that were not measured. However, the consequences of adaptation to both temperature and nutrition on thermal tolerance and its plasticity were not assessed.

In this study, we aimed to understand how long-term adaptation to combined thermal-nutritional stress affects larval thermotolerance and thermal plasticity. We assessed basal and induced thermotolerance of larvae from nine *D. melanogaster* experimental evolution lines that were evolved under combinations of three temperatures (18°C, 25°C, and 28°C) and three diets (standard, diluted, and low-protein:high-carbohydrate [low P:C] diets) using laboratory natural selection (Alton *et al*. 2024). These conditions were chosen to reflect variation in the nutritional context that developing larvae experience in rotting fruit, with the diluted diet representing reduced food availability and the low P:C diet representing reduced food quality (Markow *et al*. 1999; Kristensen *et al*. 2011; Almeida de Carvalho and Mirth 2017; Silva-Soares *et al*. 2017) and to simulate future thermal and nutritional conditions in the context of long-term climate change (DaMatta *et al*. 2010; Gowik and Westhoff 2011; IPCC 2018). The diluted and low P:C diets contained the same concentration of protein, and the standard and low P:C diets contained the same caloric concentration by replacing the calories from protein with carbohydrates. This allowed us to examine the effect of protein and carbohydrate concentrations of the selection diets independently of each other.

Only larvae were exposed to selection to reflect their limited capacity to avoid stress by behaviourally choosing or moving to more favourable microhabitats in the wild (Austin and Moehring 2019). Based on previous studies examining thermal and nutritional adaptation individually, we first predicted that adaptation to 28°C would increase larval heat tolerance (Huey et al. 1991; Cavicchi et al. 1995), and this effect would be strongest for lines adapted to the standard diet, high in protein (Kolss et al. 2009).

Second, we predicted that adaptation to 18°C would result in increased cold tolerance, and that this effect would be greater for lines adapted to the low P:C diet. Finally, if basal thermotolerance trades off with thermal plasticity, we would expect plasticity to be reduced in any lines that had evolved higher basal thermotolerance.

Our results provide insight into the complexity of evolutionary responses to combinations of thermal and nutritional stress, and what this means for our ability to predict responses to ongoing climate change. Previous work in microbes reveals that adaptation to combinations of stressors can be complex and difficult to predict (Karve *et al*. 2016). Consistent with this work, our results reveal that adaptation to combinations of thermal and nutritional stress results in both genetic and plastic shifts in pre-adult traits that are not easily predicted by adaptation to single stressors. Finally, they highlight that studies that address the complexity of adaptation to multiple stressors are needed to better understand responses to ongoing environmental change.

## Methods

### Fly stock – Collection and initiation of experimental evolution lines

Two hundred female *D. melanogaster* were collected in Duranbah, Queensland (28.3° S, 153° E) in 2018 to initiate isofemale lines (Alton *et al*. 2024). Flies were returned to the laboratory and were treated with tetracycline to remove *Wolbachia* (Min and Benzer 1997). Two generations after tetracycline treatment, five males and five virgin females from each line were used to create a mass-bred base population. A large number of isofemale lines were used to maximise genetic variation in the founding mass-bred population. The base population was maintained under control conditions — at 25°C on the standard diet with 12:12 hour light:dark cycles (see below) — across 60 bottles each containing approximately 750–1000 flies, for two generations before the initiation of selection. It is worth noting that, because the mass-bred population was maintained for only two generations before the selection experiment, it was most likely not fully adapted to laboratory conditions and was therefore not at equilibrium. This, in turn, may have contributed to linkage disequilibrium in the base (starting) population (e.g. Kellermann *et al*. 2015). That said, we believe that the control lines at least would have reached equilibrium by the time of the experiments described in this study, given laboratory adaptation occurs relatively quickly, and is expected to have stabilised by 7–10 generations (Harshman and Hoffmann 2000; Hoffmann *et al*. 2001). Because we used laboratory natural selection (rather than directional selection), it is also likely that the selected lines had reached equilibrium by the time the experiments were conducted (Sgrò and Blows 2004).

### Experimental evolution regimes and selection protocol

Eggs were collected from the base population and were divided among nine combined thermal-nutritional selection treatments with five replicate lines per treatment (Alton *et al*. 2024). These treatments consisted of three temperatures (18°C, 25°C, and 28°C) and three diets (standard, diluted, and low P:C) (Table S1) under a 12:12 light:dark cycle. Only larvae were exposed to selection. All diets contained varying amounts of potato flake, inactive yeast, agar, and dextrose, with the addition of nipagin and propionic acid to prevent bacterial and fungal growth (Holleley *et al*. 2008) (Table S1).

The standard diet had a 1:3 protein-to-carbohydrate (P:C) ratio (yeast 36.36 g/L) and contained 320 kcal/L; the diluted diet had a 1:3 P:C ratio (yeast 9.09 g/L) and contained 80 kcal/L; and the low P:C diet had a 1:12 P:C ratio (yeast 8.18 g/L) and contained 320 kcal/L (Table S1).

These selection conditions reflect combinations of thermal and nutritional changes that may be expected in the future under climate change; 18°C and 25°C represent the current average Australian winter and summer temperatures respectively (Alton *et al*. 2024), while 28°C represents a 3°C increase of the average summer temperature projected in twenty years (Australian Academy of Science 2021). The diluted diet and the low P:C diet simulate a reduction of protein and varying carbohydrates in plant composition projected under climate change (DaMatta *et al*. 2010), and span the range of nutritional contexts experienced by developing larvae in rotting fruit (Matavelli *et al*. 2015; Silva-Soares *et al*. 2017). The 25°C standard diet selection regime was used as a control and a reference point for further analysis.

At the start of each generation of selection, each selection replicate line was established with three bottles containing 200–250 eggs on 70 mL of selection diets to equalize density across all selection regimes. Animals were allowed to develop on their respective diet and temperatures until adult eclosion. We avoided inadvertent selection on development time by collecting all eclosing adults over a 5-day period across selection lines (all adults had emerged over a 5-day period, even those lines that took the longest to develop). Adults from the same replicate line were mixed and separated into two bottles containing the standard diet at 25°C for three days to allow mating. On the fourth day, adults were transferred to 250 mL egg-laying chambers containing 11 mL of egg-laying medium, the standard diet with double the concentration of agar of the standard fly diet and a layer of autoclaved yeast for acclimation. We doubled the concentration of agar to prevent females from embedding their eggs into the food, making egg collection easier. After the 24-hour acclimation, fresh medium was provided for egg-laying overnight. The eggs were collected the next morning to establish the next generation as described above. This means that adults were between 5–10 days old at the time of egg collection, and we may therefore have inadvertently selected for early life fecundity.

At the time of the experiment described below, generations of selection were as follows: 61 for the 18°C standard diet, 51 for the 18°C diluted and low P:C diets, 89 for the 25°C standard diet, 75 for the 25°C diluted and low P:C diets, 100 for the 28°C standard diet, 81 for the 28°C diluted diet, and 84 for the 28°C low P:C diet. Due to logistical limitations, three replicate lines were randomly selected from the five set up for each selection regime to perform the experiment described below. Prior to the experiment, all lines were placed in common garden conditions at 25°C on a standard diet for at least two generations to minimise parental effects (De Villemereuil *et al*. 2016).

### Plastic and evolved shifts in larval thermotolerance Experimental set up

After two generations of common garden rearing, adults were transferred to laying plates containing egg-laying medium and a surface layer of autoclaved yeast to induce oviposition. Laying plates were placed at 25°C for 14 hours to allow egg-laying. Eggs were collected from laying plates and transferred to vials containing 7.2 mL of standard diet, at a density of twenty eggs per vial (Agnew *et al*. 2002). Five replicate vials were set up for each replicate line per selection regime. First instar larvae (hatched 24 hours after egg transfer) were exposed to the hardening and acute stress treatments. Vials were maintained at 25°C under 12:12 hour light:dark cycles throughout the experiment, except when exposing larvae to the hardening and acute stress treatments described below. This common garden experimental design enabled us to investigate the effects of genetic adaptation to combined thermal-nutritional stress on larval thermotolerance and thermal plasticity.

### Assessing induced (plastic) and basal thermotolerance

Larval cold plasticity was assessed by subjecting 24-hour-old first instar larvae, reared from embryos at 25°C on a standard diet, to a two-hour hardening treatment in a 0°C water bath, followed by a 24-hour recovery period at 25°C. After recovery, these 48-hour-old larvae were exposed to a four-hour acute cold shock at 0°C in a water bath and then returned to 25°C under 12:12 hour light: dark cycles to complete development. Basal cold tolerance was assessed by exposing 48-hour-old larvae, reared from embryos at 25°C on a standard diet, to a four-hour acute cold shock at 0°C in a water bath, without prior hardening treatment. The larvae were then allowed to complete development at 25°C under 12:12 hour light:dark cycles. 48-hour-old larvae were exposed to the acute cold shock treatment to equalise the stage at which larvae were exposed to the acute stress across the hardened and basal treatments.

Larval heat plasticity was assessed by subjecting 24-hour-old first instar larvae, again reared from embryos at 25°C on a standard diet, to a one-hour hardening treatment in a 35°C water bath, followed by a 24-hour recovery period at 25°C. After recovery, these 48-hour-old larvae were exposed to an acute heat shock of 39°C for 30 minutes. Those larvae were then allowed to develop at 25°C under 12:12 hour light:dark cycles until adult eclosion. Basal heat tolerance was assessed by exposing 48-hour-old larvae, reared from embryos on the standard diet at 25°C, to an acute heat shock treatment of 39°C for 30 minutes without any hardening treatment. They were then allowed to complete development at 25°C under 12:12 hour light:dark cycles.

A control treatment was also included, where vials contained larvae that were not exposed to any heat or cold stress treatments and were maintained on control food at 25°C under 12:12 hour light:dark cycles until adult eclosion.

The temperatures and durations for both heat and cold shock hardening and acute treatments were determined by pilot experiments (details in supplementary materials). Throughout these treatments, all vials were temporarily sealed with plastic caps and parafilm to prevent water leakage. After adult eclosion, larval-adult viability was subsequently measured as the ratio of the number of emerged adult flies to the initial egg number deposited in the vials.

### Statistical analyses

All data analyses and visualisations were performed in R (R Core Team 2024). The larval-adult viability data from all treatments were fitted with generalised linear mixed-effects models with a binomial distribution using the *lme4* package (Bates *et al*. 2015). All model weights were set to the number of eggs deposited per vial (20 eggs).

To test for the effects of long-term selection under combined thermal-nutritional stress on larval-adult viability under control (25°C standard diet) conditions, selection treatments (combinations of selection temperature and diet) were included as a fixed effect, while block (the day that eggs were picked to set up the experiment) and replicate line were included as non-nested (crossed) random effects to account for variation between blocks or variation among replicate lines; these random effects were included to improve model accuracy but were not tested for statistical significance (Bates *et al*. 2015).

To test for the effects of selection on larval basal thermotolerance and thermal plasticity, selection temperature, selection diet, hardening treatment (hardened and non-hardened), and their three-way interaction were included as fixed effects. Block and replicate line were treated as non-nested (crossed) random effects to account for any effect of block or variation among replicate lines; these random effects were not tested for statistical significance (Bates *et al*. 2015). Interactions between the random and fixed effects were also not tested because they were not core to our question.

Significance of the fixed effects was assessed using ANOVA using the *car* package (Fox *et al*. 2001), which uses the Wald Chi-square test to generate p-values for the fixed effects (Bolker *et al*. 2009; Rebolledo *et al*. 2023). Post hoc multiple comparisons were performed using the *emmean* package (Lenth 2023) to test for significant differences in larval-adult viability across levels of each factor. To quantify the amount of variation in the data explained by our models, we estimated conditional R2 values which take both fixed and random effects in the model into account (Nakagawa *et al*. 2017) using the *MuMIn* package (Nakagawa and Schielzeth 2013), since traditional measures of effect size are not applicable to generalised linear mixed models (Nakagawa *et al*. 2017).

## Results

Our aim was to understand the effects of long-term adaptation to combined thermal-nutritional stress on larval-adult viability, thermotolerance, and thermal plasticity. To address this, we first asked whether adaptation to combinations of three temperatures (18°C, 25°C, and 28°C) and three diets (standard, diluted and low P:C) resulted in evolved shifts in viability under control (25°C, standard diet) conditions. We then assessed basal and induced (plasticity in) cold and heat tolerance of *D. melanogaster* larvae from the evolved lines.

### Larval-adult viability under control conditions

We first explored whether there was evidence for any response to selection in larval-adult viability under control (non-stress) conditions. We compared larval-adult viability from all selection lines to our control 25°C standard diet line, and found that selection treatments had no effect on larval-adult viability (conditional R2 = 0.14, Table 1, Figure 1). This implies that adaptation to combinations of temperature and diet did not incur costs in the form of reductions in viability under control conditions.

**Figure 1.**
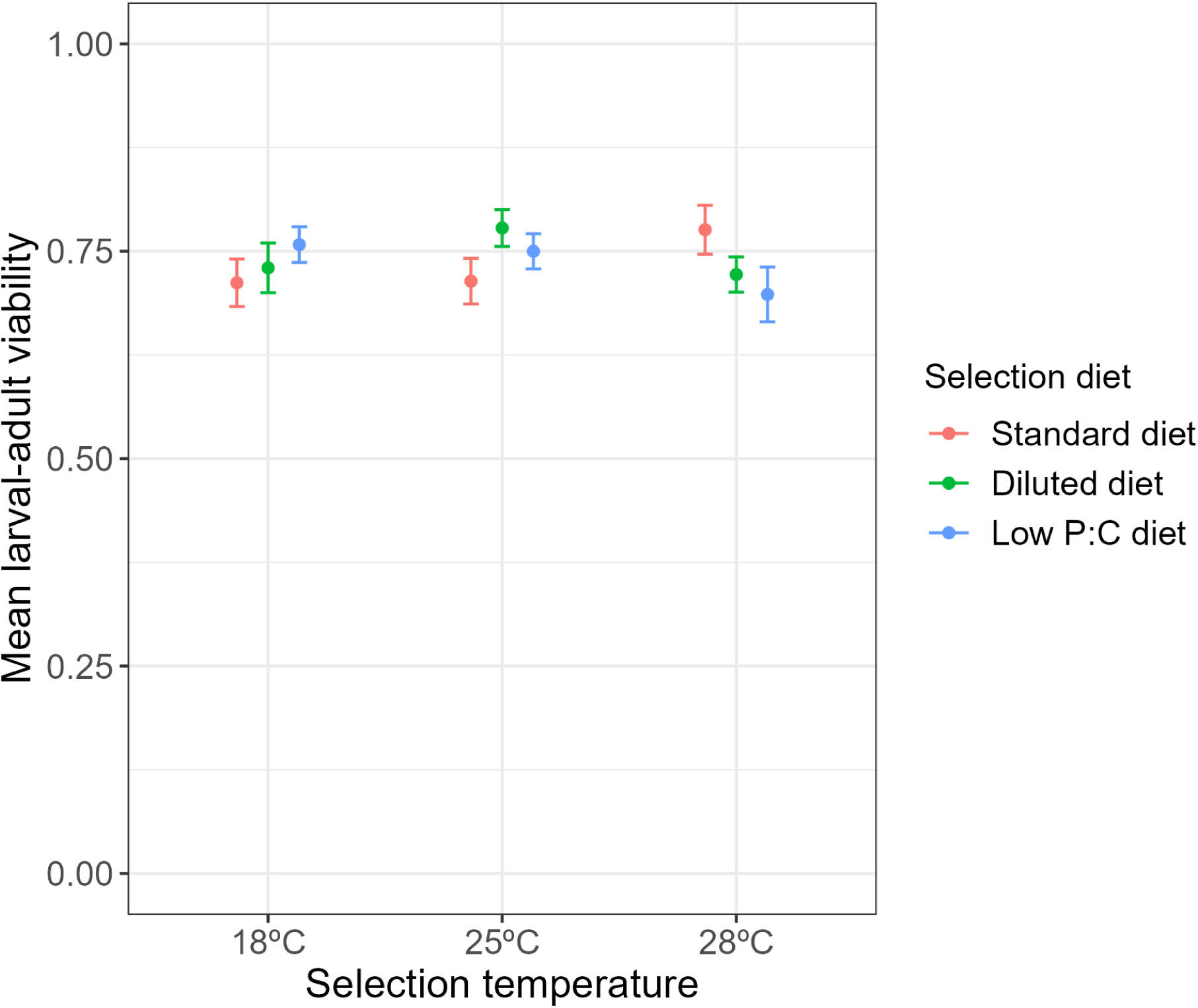
Mean larval-adult viability of experimental evolution lines tested under control treatments. Control treatment involved larvae developing on standard (control) food at 25°C under 12:12 light:dark cycles until adult eclosion. Error bars indicate ± 1 standard error.

**Table 1.** Results of multiple comparisons on the effects of selection treatment on larval-adult viability under control (no stress) conditions. Significance is defined as a p-value less than 0.05.

| Contrast: Selection treatment | Estimate | SE | Z ratio | P-value |
| --- | --- | --- | --- | --- |
| 18°C standard diet – 25°C standard diet | -0.030 | 0.142 | -0.211 | 1.000 |
| 18°C diluted diet – 25°C standard diet | 0.041 | 0.143 | 0.285 | 1.000 |
| 18°C low P:C diet – 25°C standard diet | 0.230 | 0.144 | 1.585 | 0.813 |
| 25°C standard diet – 25°C diluted diet | -0.357 | 0.148 | -2.414 | 0.276 |
| 25°C standard diet – 25°C low P:C diet | -0.194 | 0.145 | -1.342 | 0.919 |
| 25°C standard diet – 28°C standard diet | -0.337 | 0.148 | -2.280 | 0.354 |
| 25°C standard diet – 28°C diluted diet | -0.056 | 0.143 | -0.390 | 1.000 |
| 25°C standard diet – 28°C low P:C diet | 0.048 | 0.141 | 0.337 | 1.000 |

### Basal cold tolerance

We then explored whether adaptation to combined thermal-nutritional stress altered basal cold tolerance. We found a significant selection diet effect (conditional R2 = 0.33, Table 2) and a significant two-way interaction between selection temperature and selection diet on basal cold tolerance (Table 2, Figure 2), meaning that any response to selection diet further changes with the selection temperature. This interaction is illustrated by the fact that lines selected on both diluted and low P:C diets had higher basal cold tolerance than those selected on standard diets but only when they had evolved at 18°C and 28°C, and not 25°C (Figure 2, Table S2). Because the diluted and low P:C diets had the same concentration of protein, this suggests that the interaction between protein concentration and temperature is driving these effects irrespective of carbohydrate levels.

**Figure 2.**
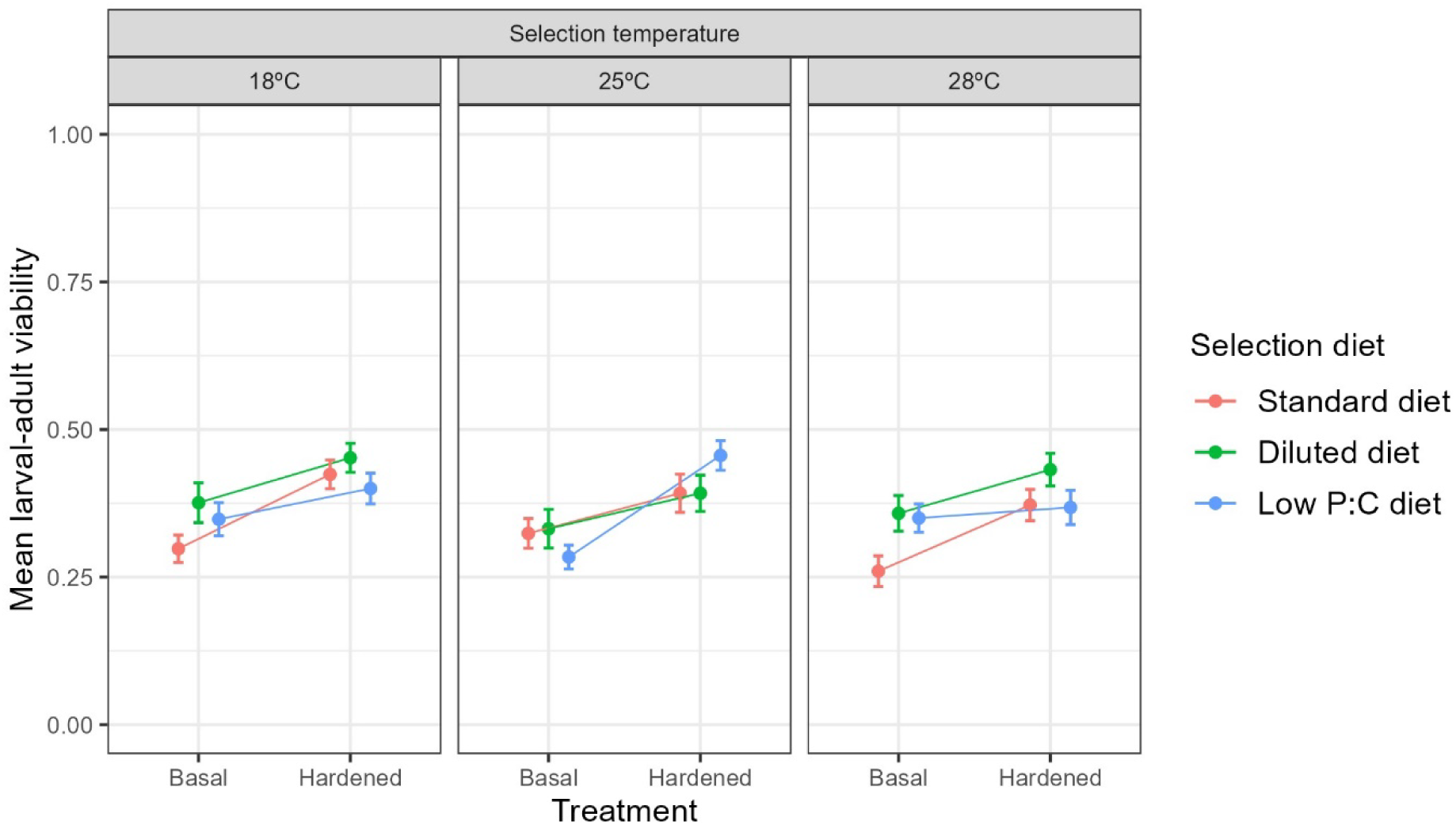
Mean larval-adult viability of basal (non-hardened) and hardened (2 h at 0°C) experimental evolution lines after exposure to an acute cold shock treatment of 4 h at 0°C. Error bars indicate ± 1 standard error.

**Table 2.**
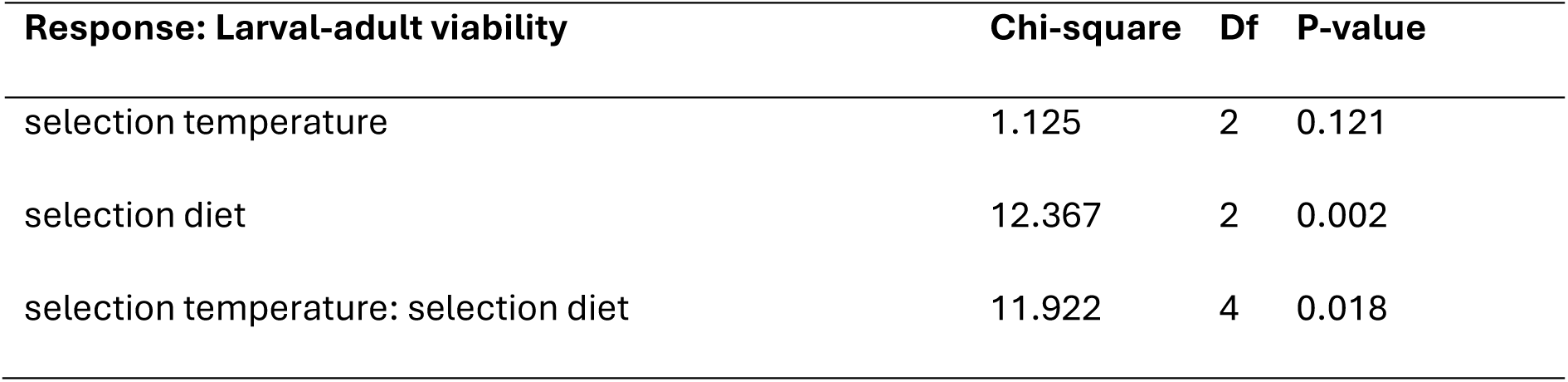
Results of analysis of variance testing for effects of selection temperature, selection diet, and their interaction on larval basal (non-hardened) cold tolerance. Significance is defined as a p-value less than 0.05.

### Plasticity in cold tolerance (cold hardening capacity)

Larval cold tolerance was significantly affected by selection diet, cold hardening treatment, and a three-way interaction between selection temperature, selection diet, and cold hardening treatment (conditional R2 = 0.26, Table 3, Figure 2). This means that the larvae showed a plastic shift in cold tolerance (a significant hardening response) but that this plastic shift was dependent on the selection diet and selection temperature. Hardening induced an increase in cold tolerance in larvae from most selection treatments, but not for those selected on the low P:C diet at 18°C and 28°C and those selected on the standard diet at 25°C (Figure 2, Table S3). This means that selection on diluted diet increased plasticity in cold tolerance in general, while selection on low P:C diet increased plasticity only at the optimal temperature, and selection on standard diet (higher protein) increased plasticity in cold tolerance only at non-optimal temperatures.

**Table 3.** Results of analysis of variance testing for effects of selection temperature, selection diet, cold hardening treatment, and their interaction on larval cold tolerance. Significance is defined as a p-value less than 0.05.

| Response: Larval-adult viability | Chi-square | Df | P-value |
| --- | --- | --- | --- |
| selection temperature | 4.677 | 2 | 0.096 |
| selection diet | 15.226 | 2 | <0.001 |
| hardening treatment | 70.606 | 1 | <0.001 |
| selection temperature: selection diet | 6.969 | 4 | 0.138 |
| selection temperature: hardening treatment | 1.948 | 2 | 0.378 |
| selection diet: hardening treatment | 1.498 | 2 | 0.473 |
| selection temperature: selection diet: hardening treatment | 16.389 | 4 | 0.003 |

### Basal heat tolerance

We then explored the effects of long-term adaptation to combined thermal-nutritional stress on basal heat tolerance. Basal heat tolerance was significantly affected by the selection temperature, but not selection diet (conditional R2 = 0.12, Table 4, Figure 3). The 25°C selection lines had higher basal heat tolerance than the 28°C, but no difference was found in basal heat tolerance between the 18°C and 25°C or between the 18°C and 28°C selection lines (Figure S1, Table S4). This suggests that long-term selection at higher temperatures does not necessarily lead to an increase in basal heat tolerance, contrary to previous studies (Cavicchi *et al*. 1995).

**Figure 3.**
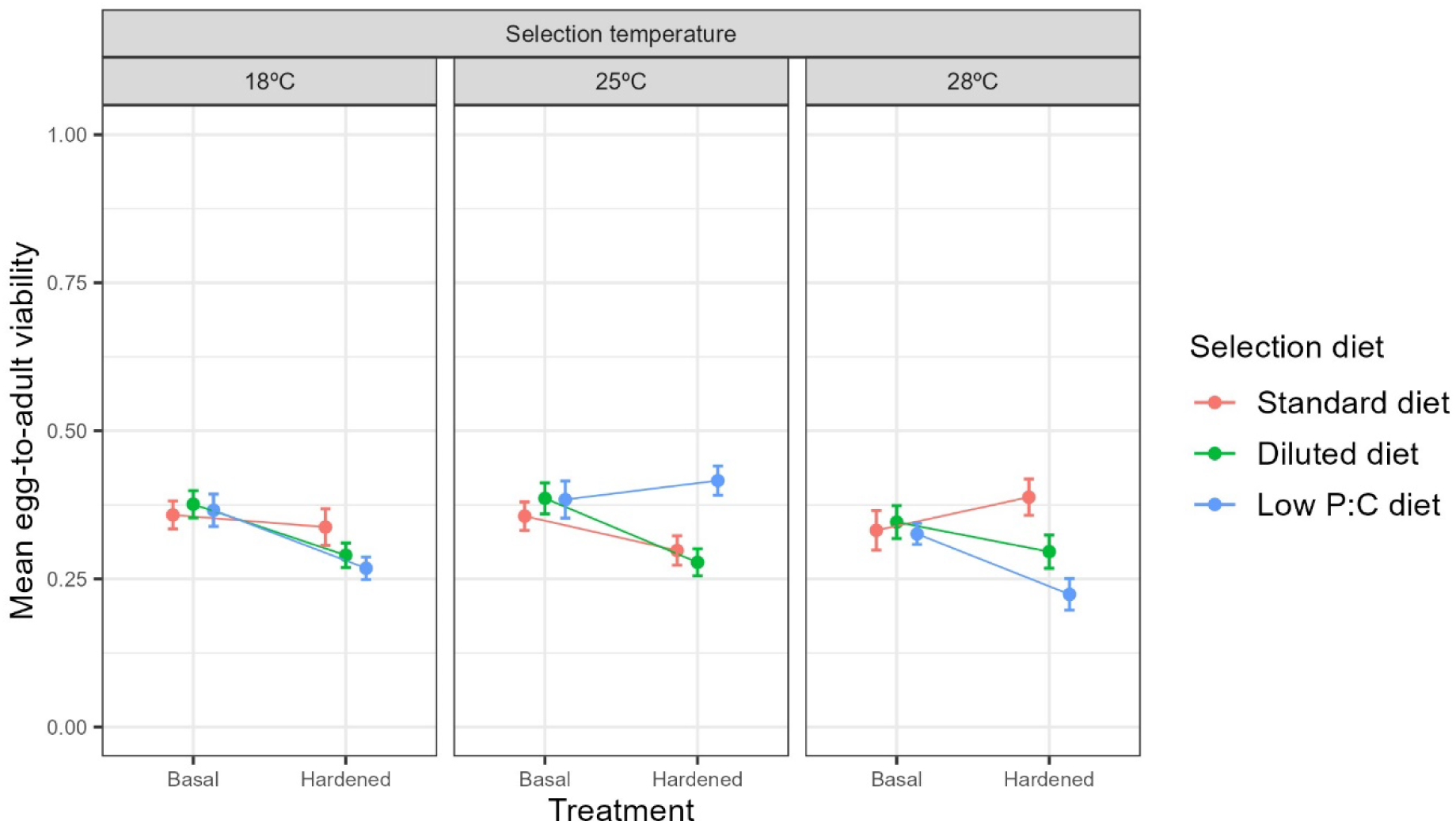
Mean larval-adult viability of basal (non-hardened) and hardened (1 h at 35°C) experimental evolution lines after exposure to an acute heat shock treatment of 0.5 h at 39°C. Error bars indicate ± 1 standard error.

**Table 4.** Results of analysis of variance testing for effects of selection temperature, selection diet, and their interaction on larval basal (non-hardened) heat tolerance. Significance is defined as a p-value less than 0.05.

| **Response: Larval-adult viability** | **Chi-square** | **Df** | **P-value** |
| --- | --- | --- | --- |
| selection temperature | 6.142 | 2 | 0.046 |
| selection diet | 2.038 | 2 | 0.361 |
| selection temperature: selection diet | 1.079 | 4 | 0.898 |

### Plasticity in heat tolerance (heat hardening capacity)

Lastly, we explored the effects of long-term adaptation to combined thermal-nutritional stress on plasticity in heat tolerance. We found a three-way interaction between selection temperature, selection diet, and hardening treatment (conditional R2 = 0.21, Table 5, Table S5). This means that plastic shifts in heat tolerance depended on both the selected diet and selected temperature.

**Table 5.** Results of analysis of variance testing for effects of selection temperature, selection diet, heat hardening treatment, and their interaction on larval heat tolerance. Significance is defined as a p-value less than 0.05.

| **Response: Larval-adult viability** | **Chi-square** | **Df** | **P-value** |
| --- | --- | --- | --- |
| selection temperature | 7.072 | 2 | 0.029 |
| selection diet | 2.557 | 2 | 0.278 |
| hardening treatment | 25.277 | 1 | <0.001 |
| selection temperature: selection diet | 31.516 | 4 | <0.001 |
| selection temperature: hardening treatment | 1.824 | 2 | 0.402 |
| selection diet: hardening treatment | 9.289 | 2 | 0.010 |
| selection temperature: selection diet: hardening treatment | 21.281 | 4 | <0.001 |

In contrast to the positive cold hardening response described above, where hardening increased cold resistance in most selection lines (Figure 2, Table S3), heat hardening resulted in no change or a reduction in heat tolerance (Figure 3, Table S5), and these effects varied across selection diet and selection temperature. Lines selected at 25°C on low P:C diets and at 28°C on standard diets, which showed no significant response to hardening (Figure 3, Table S5); however, heat hardening resulted in a significant reduction in heat tolerance in all other lines (Figure 3, Table S5).

## Discussion

Temperature and nutrition are amongst the most common stressors experienced by animals in nature, and this will become increasingly so with ongoing climate change (Rosenblatt and Schmitz 2016). However, assessments of climate risk have largely focused on the ability to tolerate thermal extremes (Hoffmann *et al*. 2003; Bowler 2005; Loeschcke and Sørensen 2005; Teets *et al*. 2020; Ma *et al*. 2021), rather than sensitivity to combinations of stressors that organisms will increasingly experience in nature.

Genetic adaptation will play a key role in the long-term population persistence (Potvin and Tousignant 1996; Gienapp *et al*. 2008; Lande 2009). However, we know very little about the extent to which populations harbour the genetic variation needed to adapt to combined thermal-nutritional stress, and whether such adaptation may be constrained by trade-offs. We also have little insight into the consequences of adaptation to both temperature and nutrition on the ability to shift phenotypes via plasticity, and whether shifts in responses can be predicted by adaptation to a single stressor alone. We addressed these outstanding questions using lines of *D. melanogaster* whose larvae evolved under combinations of three temperatures (18°C, 25°C, and 28°C) and three diets (standard, diluted, and low P:C) (Alton *et al*. 2024), and assessed the extent to which adaptation to these stressors resulted in reduced viability or shifts in larval thermotolerance and thermal plasticity.

### Temperature and nutrition interact to affect evolutionary shifts in pre-adult viability of D. melanogaster under control conditions

Previous studies have shown that populations harbour the genetic variation needed to adapt to diet (Kolss *et al*. 2009; Kristensen *et al*. 2011) and temperature (Sørensen *et al*. 2001; Kimura 2004; Sgrò *et al*. 2010; Freda *et al*. 2019) individually and in combination (Bochdanovits and De Jong 2003), which is consistent with the current study. However, these previous studies provide mixed and context-dependent evidence for trade-offs associated with adaptation. For example, adaptation to changes in nutrition alone can result in reduced (Kristensen *et al*. 2011) or increased (Leftwich *et al*. 2017) pre-adult viability regardless of the test diet, or an increase in viability only when tested on the evolved diet (Kolss *et al*. 2009). Similarly, thermal evolution can result in increased pre-adult survival only when tested at the temperature at which populations evolved (Partridge *et al*. 1994; Santos *et al*. 2006).

Such complexity is also evident when larvae are forced to adapt to both temperature and diet simultaneously. Specifically, Bochdanovits and De Jong (2003) found that adaptation to poor larval food in *D. melanogaster* resulted in increased viability when lines were selected at 27.5°C but decreased viability when lines were selected at 17.5°C. In contrast, we found that adaptation to low-protein diets (the diluted and low P:C diets) reduced larval viability compared to larvae evolved on standard food when selected at 28°C, whereas this pattern was largely reversed in lines evolving at 25°C. Lines evolved on different diets did not differ in their larval viability when evolved at 18°C. Our results suggest that simultaneous adaptation to nutritional stress and elevated temperature may negatively impact population growth, at least when reared under control (non-stress) conditions. Studies testing evolved shifts in viability and trade-offs under a range of conditions, such as using a reciprocal design, are needed to better understand the consequences for population growth.

### Adaptation to temperature and nutrition in combination affects larval thermotolerance and thermal plasticity

Previous studies on adult thermal adaptation (Huey *et al*. 1991; Cavicchi *et al*. 1995; Bubliy and Loeschcke 2005) and dietary plasticity (Hazel 1995; Kostal and Simek 1998; Andersen *et al*. 2010; Colinet *et al*. 2013) suggest that adaptation to lower temperatures or a high-carbohydrate diet increases basal cold tolerance. We predicted that lines exposed to 18°C and the low P:C diet (high carbohydrate) would evolve higher basal cold tolerance compared to lines from other selection temperatures and diets, although this assumption ignores complex interactions between temperature and diet, and is based on studies focussed largely on adult traits. Surprisingly, lines selected on poor diets, both diluted and low P:C, evolved higher basal cold tolerance compared to lines selected on a standard diet, but only when they were selected at the non-optimal temperatures of 18°C and 28°C. This contradicts previous single nutritional stressor studies (Hazel 1995; Kostal and Simek 1998; Andersen *et al*. 2010; Colinet *et al*. 2013) that suggest a high-carbohydrate diet increases cold tolerance, except that Kolss et al. (2009) were consistent with our findings and suggested that adaptation to poor (25% diluted) food at 25°C had no effect on basal cold tolerance. We found that cold tolerance may be enhanced when populations are forced to adapt to combinations of non-optimal thermal and low-protein nutritional conditions, as the diluted and low P:C diets were protein-matched. This further emphasises that evolutionary responses to combined stressors cannot be predicted by single-stressor studies alone. More studies are needed to better understand the effects of a broader range of temperature and nutritional contexts on basal cold tolerance and the underlying mechanisms.

While some studies suggest trade-offs between basal cold tolerance and cold plasticity (Bubliy and Loeschcke 2005; Nyamukondiwa *et al*. 2011; Noh *et al*. 2017), we found that most experimental evolution lines showed a positive cold hardening response—cold tolerance increased with hardening. This suggests that basal cold tolerance and plasticity in cold tolerance may have evolved together, depending on the environmental conditions to which populations are exposed. Further studies using different rearing conditions are recommended to further assess the trade-offs between basal cold tolerance and cold plasticity.

Adaptation to elevated temperatures (Cavicchi *et al*. 1995; Bettencourt *et al*. 1999) and plastic responses to high-protein diets (Andersen *et al*. 2010) have been shown to increase adult heat tolerance. We therefore predicted that selection at 28°C on the standard (high-protein) diet would increase larval heat tolerance. However, we found that larvae selected at 25°C had the highest average basal heat tolerance. This discrepancy between our results (a significant effect of selection temperature but not diet) and previous studies (Andersen *et al*. 2010, which reported a significant effect of selection diet) may be due to the fact that previous studies focused on adult, not larval, thermotolerance, and insect life stages can differ in their thermal sensitivity (Kingsolver and Buckley 2020). For instance, thermal stress tolerance in *D. buzzatii* was either independent or weakly correlated across life stages (Sarup *et al*. 2006). Similarly, different life stages of *D. melanogaster* also vary in their basal heat tolerance (Freda *et al*. 2019; Moghadam *et al*. 2019), although the precise mechanisms controlling this variation remain unclear (Bowler and Terblanche 2008).

We also found that plasticity in heat tolerance became maladaptive after adaptation to most of the temperature-diet selection treatments. Specifically, the hardening treatment seemed to impose further heat stress on larvae and reduced, rather than increased, their survival and heat tolerance. Our results are partly consistent with previous work by Cavicchi *et al*. (1995), who found increased heat tolerance after hardening in lines adapted to 18°C and 25°C, while lines adapted to 28°C lost the ability to mount a hardening response. Additionally, the 28°C lines showed increased sensitivity to higher hardening temperatures (43°C), leading to a decrease in their hardened heat tolerance (Cavicchi *et al*. 1995). In contrast, we found that none of the evolved lines showed a significant positive hardening response; all significant hardening responses reduced larval heat tolerance.

Our results could be explained by plasticity in heat tolerance reaching a physiological limit. We found very weak evidence for evolved shifts in basal heat tolerance, suggesting that the starting population may have had limited genetic variation for larval heat tolerance (Loeschcke and Krebs 1996; Lansing *et al*. 2000) and its ability to evolve plasticity. Limited responses to selection on larval heat tolerance also suggest that a trade-off between basal and hardened plasticity (Gunderson and Stillman 2015; van Heerwaarden *et al*. 2016; van Heerwaarden and Kellermann 2020) is unlikely to explain our findings.

Another explanation could be that the threshold for inducing a heat hardening response might have shifted down as a consequence of long-term adaptation to combined thermal-nutritional stress in the present study (Cavicchi *et al*. 1995; van Heerwaarden *et al*. 2024). Cavicchi *et al*. (1995) found that lines adapted to 18°C had a lower threshold for inducing a hardening response compared to those adapted to 25°C. In contrast, they found that lines adapted to 28°C appeared to lose the capacity to mount a hardening response at lower hardening temperatures and exhibited a negative hardening response at the highest temperature. This is in part consistent with our results.

Plasticity of larval heat tolerance has been shown to vary across assay temperature and duration of exposure, and is species-specific (Chen *et al*. 1991; Moghadam *et al*. 2019; van Heerwaarden *et al*. 2024). Chen *et al*. (1991) demonstrated that the duration of hardening treatment significantly affected adult emergence of *Sarcophaga crassipalpis* after a heat shock; adult emergence was maximised with 2 hours of hardening at 40°C, but only achieved 50% with a 1-hour hardening treatment. Moghadam et al., (2019) further showed that heat hardening capacity depends on the temperature of acute heat stress. Hardening treatments had a negative effect on subsequent heat tolerance in 3rd instar larvae of *D. melanogaster* when the acute stress temperature exceeded 40°C, but not when it was below 40°C. Our choice of temperature and duration for hardening and acute heat stress treatments might have been too stressful for the selected larvae.

In our pilot study (see supplementary materials), larvae selected on the standard diet across different selection temperatures showed positive hardening responses for heat tolerance (Figure S3, albeit the effect was small). This means that positive hardening responses may be observed if a lower acute stress temperature was used. Future studies assessing evolutionary responses of thermotolerance and thermal plasticity should explore a broader range of temperatures, nutritional contexts, hardening treatments/durations under a reciprocal design, and across different life stages to better understand the complexity of adaptation to environmental change.

Finally, we acknowledge that the mass-bred population was maintained for only two generations before the selection experiment was initiated, meaning it was likely not fully adapted to the laboratory conditions, and therefore not at equilibrium (Kellermann *et al*. 2015). This raises the possibility that linkage disequilibrium persisted in our lines during the first 4–5 generations of selection, which may have influenced the results. Nonetheless, we believe this study profvides important insight into the complexities of adaptation to combinations of stressors.

In conclusion, we found that long-term adaptation to low-protein diets and elevated temperature simultaneously resulted in reduced larval viability, at least under control conditions. Further, we observed unpredictable shifts in larval basal thermotolerance and its plasticity as a result of adaptation to combined thermal-nutritional stress, which could not be predicted from studies of thermal or nutritional adaptation individually.

We found that basal cold tolerance and plasticity in cold tolerance may co-evolve. However, the evolved lines seemed to lose the ability to increase heat tolerance via positive heat hardening responses, and largely displayed reductions in heat tolerance following hardening. Taken together, our results suggest that being forced to adapt to shifts in both the thermal and nutritional environments simultaneously may increase the vulnerability of ectotherms to ongoing climate change.

## Results – table and graphs

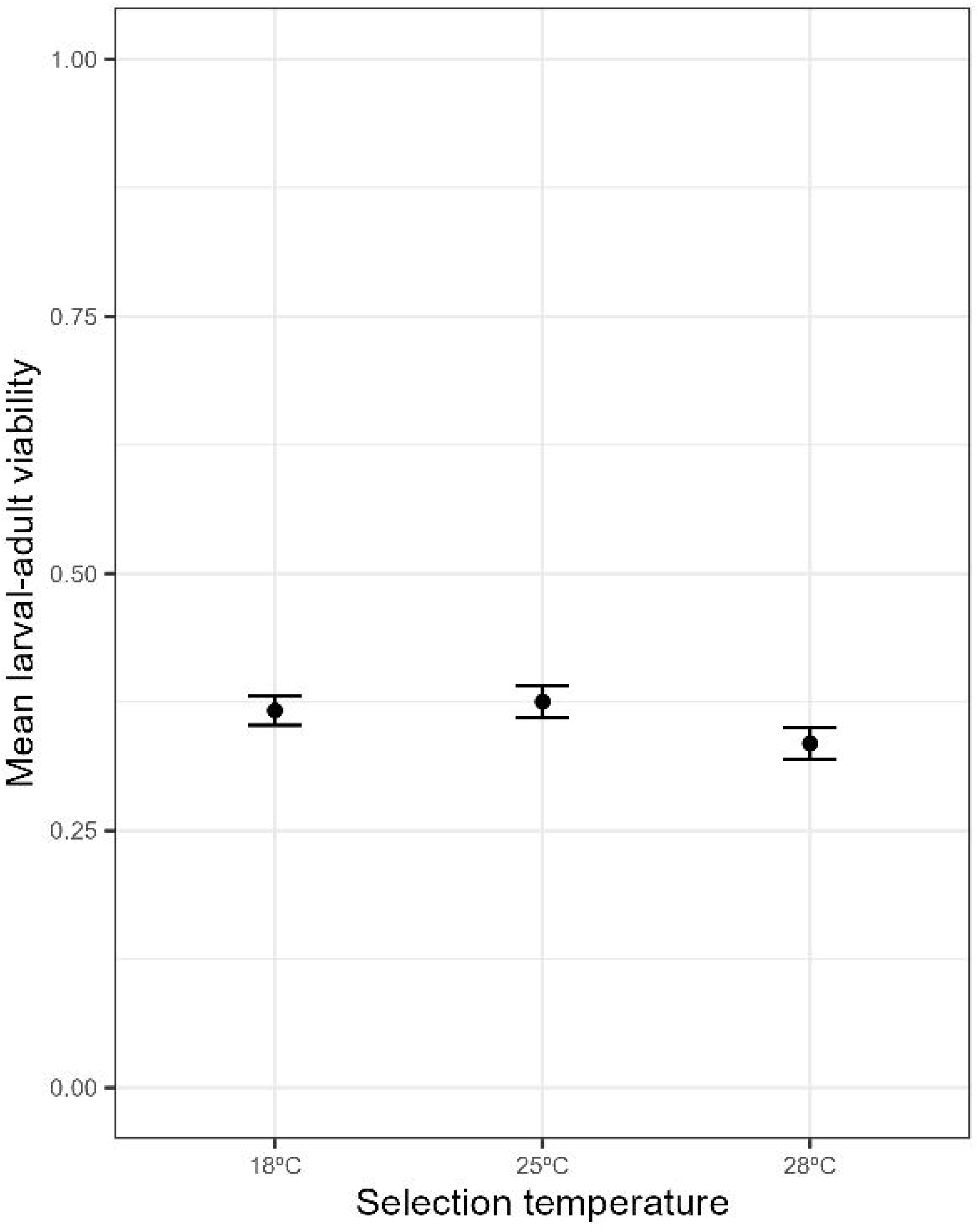

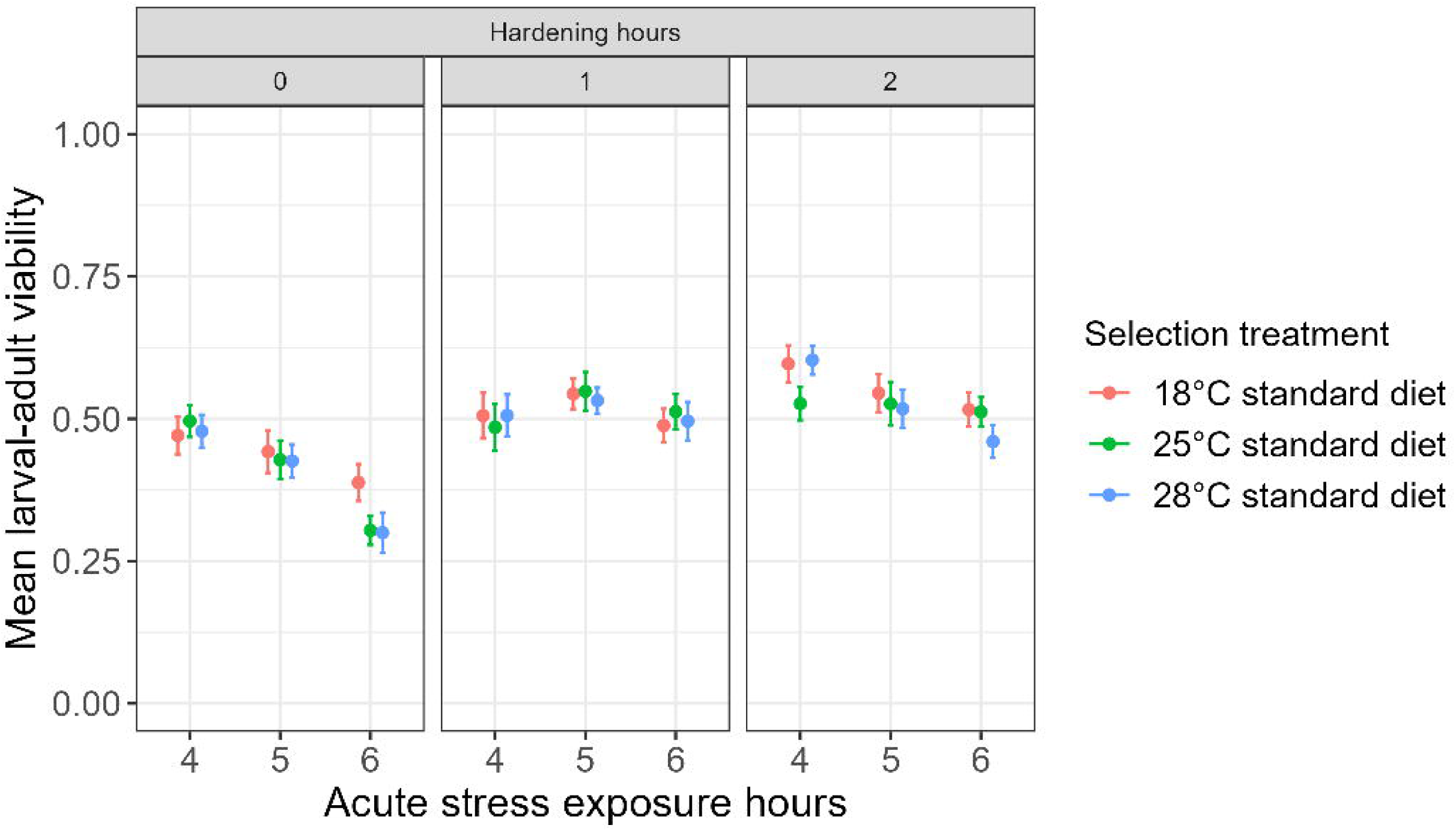

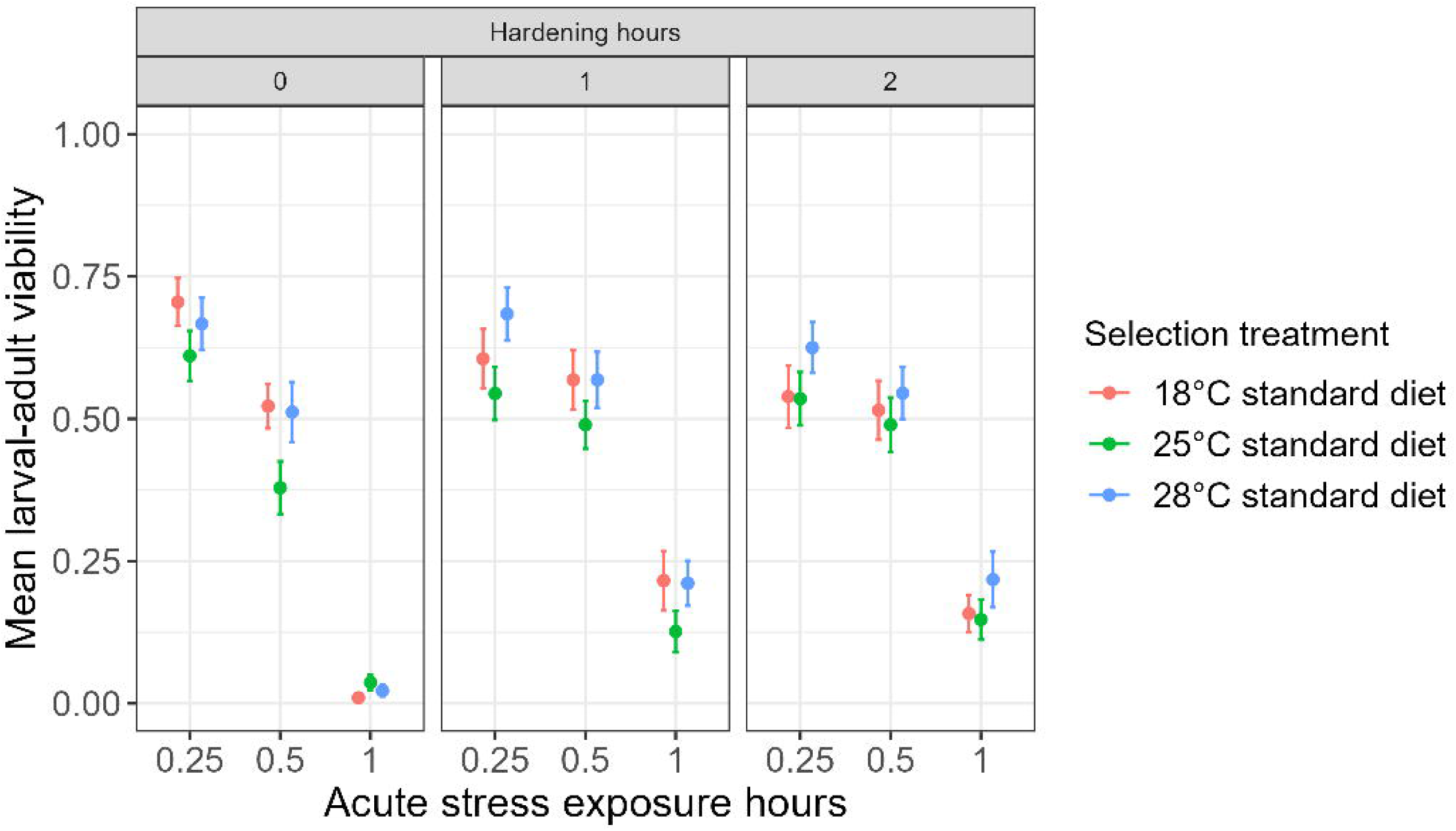

