## Supplementary material for "Assessing the effect of experimental evolution under combined thermal-nutritional stress on larval thermotolerance and thermal plasticity in *Drosophila melanogaster*": Table S1

### Methods

*Pilot experiment – determining temperatures and duration of exposure for hardening and acute thermal stress treatments*

Only lines that were selected on standard diets were used in pilots due to logistical limitations. After at least two generations of common garden rearing, adults from the experimental evolution lines were transferred to laying plates containing the standard diet with doubled agar. A layer of autoclaved yeast was painted on the food surface to induce oviposition. Laying plates were placed at 25°C for 14 hours to allow egg laying. Eggs were collected from laying plates and transferred to vials containing 7.2 mL standard diets, with a density of twenty eggs per vial (Agnew *et al.* 2002). Five replicate vials were set up for each replicate line for each selection regime. Vials were maintained at 25°C under 12:12 hour light:dark cycles throughout the experiment, except when exposing larvae to the hardening and acute thermal stress treatments described below.

To determine conditions for the cold hardening and acute treatments, first instar larvae were subjected to either no hardening treatment (maintained at 25°C with a 12:12 hour light:dark cycle), hardening treatments at 0°C for one hour or two hours. After hardening, larvae were recovered at 25°C for 24 hours under a 12:12 hour light:dark cycle. They were then exposed to one of the four-, five-, and six-hour acute cold stress treatments at 0°C. After the acute cold stress treatments, larvae were allowed to develop at 25°C under 12:12 hour light:dark cycles until adult eclosion. All vials containing larvae were temporarily sealed with a plastic cap and parafilm to prevent water leakage during hardening and acute cold stress exposure treatments.

A similar protocol was used to determine conditions for the heat hardening and acute heat stress treatment. Hardening treatments included either no hardening treatment, 1-hour or 2-hour hardening treatments at 35°C, followed by a 24-hour recovery at 25°C under a 12:12 hour light:dark cycle. Initially, acute heat stress treatments included exposure to 39°C for either 0.5, 1, or 2 hours. As no larvae survived in the 2-hour exposure to acute heat stress in the first trial, exposure time was reduced from 2-hour to 0.25-hour. Larval-adult viability was subsequently measured as the ratio of number of adults that emerged to the initial number of eggs deposited into the vials.

*Pilot experiment - Statistical analysis*

Larval-adult viability was fitted using generalised linear mixed-effect models with a binomial distribution as its values were within the range of 0 to 1. Analysis of variance (ANOVA) was performed using lme4 package (Bates *et al.* 2015) in R (R Core Team 2024) to test for the effects of different combinations of hardening hours and acute stress exposure hours on inducing larval thermal plastic response. Selection treatment (combined temperature and diet), hardening hours, and acute stress exposure hours were treated as fixed effects, whereas block and replicate line were treated as random effects. Prior weights were set to the number of eggs deposited in one vial (i.e., 20). Multiple comparisons were performed using the 'emmeans' package (Lenth 2023) to test for the significant differences in larval-adult viability between different durations of hardening treatments and acute stress exposures.

### Results

*Cold hardening and acute stress exposure hours*

We found significant effects of hardening hours and acute stress exposure hours, and a significant two-way interaction between hardening hours and acute stress exposure hours on viability (Table S6, Figure S2). Post-hoc tests suggested 2-hour hardening treatments induced the greatest plasticity in cold tolerance in all subsequent acute stress exposures (Table S7, Figure S2). Although the extent of induced plasticity is the greatest when larvae were first exposed a 2-hour hardening treatment and further exposed to acute stress for 6 hours, followed by 5 hours and 4 hours (Table S7, Figure S2), those exposed to 4-hour acute stress consistently showed higher viability compared to those treated with 6 hours of acute stress (Table S7). We therefore set our experimental durations of cold hardening and acute cold stress exposure to 2 hours and 4 hours respectively, to account for possible lower viability in lines that were selected on diluted and low P:C diets.

*Heat hardening and acute stress exposure hours*

We found significant single effects of selection treatment, hardening hours, and acute exposure hours and a three-way interaction between these three terms (Table S8, Figure S3). Additionally, we found two-way interactions between selection treatment and hardening hours, and between hardening hours and acute stress exposure hours (Table S8). Post-hoc tests revealed that larvae subjected to 1-hour hardening followed by 0.5-hour and 1-hour acute heat exposures had significantly higher viability compared to non-hardened larvae (Table S9). Viability of non-hardened larvae with 1-hour acute stress exposure was close to zero (Figure S3). We therefore decided to set our experimental parameters to 1-hour heat hardening at 35°C and 0.5-hour acute heat stress exposure at 39°C. Interestingly, lines selected at 25°C were shown to have the greatest increase in heat tolerance after 1-hour hardening and 0.5 hours acute stress exposure, followed by 18°C and 28°C lines (Table S10).

**Table S1**. The amount of ingredients added per 1.1 litre of each diet.

| **Diet** / **Ingredients** | **Standard diet** | **Diluted diet** | **Low P:C diet** |
|---|---|---|---|
| Potato starch (g) | 20 | 5 | 20 |
| Yeast (g) | 40 | 10 | 9 |
| Agar (g) | 7 | 8 | 7 |
| Dextrose (g) | 30 | 7.5 | 55 |
| Nipagin (mL) | 12 | 12 | 12 |
| Propanoic acid (mL) | 5 | 5 | 5 |

**Table S2**. Results of multiple comparisons on the effects of the two-way interaction between selection temperature and selection diet on larval basal cold tolerance. Significance is defined as a p-value less than 0.05.

| **Selection diet = standard diet** | **Estimate** | **SE** | **Z ratio** | **P-value** |
|---|---|---|---|---|
| 18°C – 25°C | -0.267 | 0.140 | 1.901 | 0.138 |
| 18°C – 28°C | -0.046 | 0.147 | 0.313 | 0.948 |
| 25°C – 28°C | 0.221 | 0.142 | 1.558 | 0.264 |
| **Selection diet = diluted diet** | | | | |
| 18°C – 25°C | 0.179 | 0.135 | 1.332 | 0.377 |
| 18°C – 28°C | 0.657 | 0.133 | 0.493 | 0.875 |
| 25°C – 28°C | -0.114 | 0.136 | 0.835 | 0.682 |
| **Selection diet = low P:C diet** | | | | |
| 18°C – 25°C | 0.304 | 0.138 | 2.198 | 0.072 |
| 18°C – 28°C | -0.030 | 0.134 | 0.221 | 0.974 |
| 25°C – 28°C | 0.334 | 0.138 | 2.407 | 0.043 |
| **Selection Temperature = 18°C** | | | | |
| Standard diet – diluted diet | -0.464 | 0.138 | -3.375 | 0.002 |
| Standard diet – low P:C diet | -0.357 | 0.139 | -2.575 | 0.027 |
| Diluted diet – low P:C diet | 0.107 | 0.133 | 0.804 | 0.701 |
| **Selection Temperature = 25°C** | | | | |
| Standard diet – diluted diet | -0.018 | 0.137 | -0.135 | 0.990 |
| Standard diet – low P:C diet | 0.213 | 0.139 | 1.530 | 0.277 |
| Diluted diet – low P:C diet | 0.231 | 0.139 | 1.666 | 0.218 |
| **Selection Temperature = 28°C** | | | | |

| | | | | |
|---|---|---|---|---|
| Standard diet – diluted diet | -0.353 | 0.140 | -2.513 | 0.032 |
| Standard diet – low P:C diet | -0.341 | 0.140 | -2.431 | 0.040 |
| Diluted diet – low P:C diet | 0.012 | 0.134 | 0.087 | 0.996 |

**Table S3**. Results of multiple comparisons on the effects of the three-way interaction between selection temperature, selection diet, and treatment on larval-adult viability in response to acute cold stress. Significance is defined as a p-value less than 0.05.

| **Estimates (P-values)** | **Selection temperature: selection diet** | | | | | | | | |
|---|---|---|---|---|---|---|---|---|---|
| **Contrast: treatment** | **18°C: standard diet** | **18°C: diluted diet** | **18°C: low P:C diet** | **25°C: standard diet** | **25°C: diluted diet** | **25°C: low P:C diet** | **28°C: standard diet** | **28°C: diluted diet** | **28°C: low P:C diet** |
| Basal - hardened | -0.630 (<0.001) | -0.360 (0.005) | -0.221 (0.092) | -0.250 (0.061) | -0.268 (0.043) | -0.762 (<0.001) | -0.452 (0.001) | -0.307 (<0.018) | -0.095 (0.473) |
| | **Selection temperature: treatment** | | | | | | | | |
| **Contrast: selection diet** | **18°C: basal** | **25°C: basal** | **28°C: basal** | **18°C: hardened** | **25°C: hardened** | **28°C: hardened** | | | |
| Standard diet – diluted diet | -0.392 (0.011) | -0.019 (0.989) | -0.421 (0.007) | -0.122 (0.608) | -0.038 (0.955) | -0.276 (0.085) | | | |
| Standard diet – low P:C diet | -0.282 (0.098) | 0.199 (0.321) | -0.389 (0.014) | 0.127 (0.588) | -0.314 (0.042) | -0.032 (0.969) | | | |
| Diluted diet – low P:C diet | 0.11 (0.684) | 0.218 (0.254) | 0.033 (0.967) | 0.249 (0.131) | -0.276 (0.082) | 0.245 (0.143) | | | |
| | **Selection diet: treatment** | | | | | | | | |
| **Contrast: selection temperature** | **Standard diet: basal** | **Diluted diet: basal** | **Low P:C diet: basal** | **Standard diet: hardened** | **Diluted diet: hardened** | **Low P:C diet: hardened** | | | |
| 18°C – 25°C | -0.179 (0.397) | 0.194 (0.312) | 0.302 (0.070) | 0.202 (0.271) | 0.286 (0.069) | -0.239 (0.153) | | | |
| 18°C – 28°C | 0.093 (0.795) | 0.063 (0.880) | -0.014 (0.994) | 0.271 (0.095) | 0.117 (0.633) | 0.113 (0.665) | | | |
| 25°C – 28°C | 0.272 (0.129) | -0.13 (0.594) | -0.316 (0.055) | 0.07 (0.855) | -0.169 (0.390) | 0.351 (0.019) | | | |

**Table S4**. Results of multiple comparisons on the effects of selection temperature on larval-adult viability of basal larvae in response to acute heat stress. Significance is defined as a p-value less than 0.05.

| **Contrast** | **Estimate** | **SE** | **Z ratio** | **P-value** |
|---|---|---|---|---|
| **Selection temperature** | | | | |
| 18°C – 25°C | -0.036 | 0.076 | 0.478 | 0.882 |
| 18°C – 28°C | 0.146 | 0.078 | 1.882 | 0.144 |
| 25°C – 28°C | 0.1825 | 0.078 | 2.353 | 0.049 |

**Table S5**. Results of multiple comparisons on the effects of the three-way interaction between selection temperature, selection diet, and treatment on larval-adult viability in response to acute heat stress. Significance is defined as a p-value less than 0.05.

| **Estimates (P-values)** | **Selection temperature: selection diet** | | | | | | | | |
|---|---|---|---|---|---|---|---|---|---|
| **Contrast: treatment** | **18°C: standard diet** | **18°C: diluted diet** | **18°C: low P:C diet** | **25°C: standard diet** | **25°C: diluted diet** | **25°C: low P:C diet** | **28°C: standard diet** | **28°C: diluted diet** | **28°C: low P:C diet** |
| Basal - hardened | 0.085 (0.526) | 0.272 (0.045) | -0.236 (0.075) | 0.401 (0.003) | 0.467 (<0.001) | 0.228 (0.094) | 0.451 (0.001) | -0.136 (0.296) | 0.516 (<0.001) |
| | **Selection temperature: treatment** | | | | | | | | |
| **Contrast: selection diet** | **18°C: basal** | **25°C: basal** | **28°C: basal** | **18°C: hardened** | **25°C: hardened** | **28°C: hardened** | | | |
| Standard diet – diluted diet | -0.087 (0.787) | -0.109 (0.684) | -0.065 (0.977) | 0.228 (0.224) | 0.085 (0.816) | 0.399 (0.009) | | | |
| Standard diet – low P:C diet | -0.029 (0.973) | -0.112 (0.671) | 0.02 (0.988) | 0.336 (0.043) | -0.521 (0.003) | 0.772 (<0.001) | | | |
| Diluted diet – low P:C diet | 0.058 (0.900) | -0.003 (1.000) | 0.085 (0.801) | 0.108 (0.725) | -0.606 (<0.001) | 0.373 (0.028) | | | |
| | **Selection diet: treatment** | | | | | | | | |
| **Contrast: selection temperature** | **Standard diet: basal** | **Diluted diet: basal** | **Low P:C diet: basal** | **Standard diet: hardened** | **Diluted diet: hardened** | **Low P:C diet: hardened** | | | |
| 18°C – 25°C | 0.008 (0.998) | -0.015 (0.993) | -0.075 (0.833) | 0.195 (0.334) | 0.052 (0.929) | -0.663 (<0.001) | | | |
| 18°C – 28°C | 0.113 (0.676) | 0.135 (0.566) | 0.163 (0.446) | -0.209 (0.263) | -0.038 (0.959) | 0.227 (0.275) | | | |
| 25°C – 28°C | 0.106 (0.709) | 0.15 (0.498) | 0.238 (0.174) | -0.403 (0.008) | -0.09 (0.799) | 0.89 (<0.001) | | | |

**Table S6**. Results of analysis of variance testing for effects of selection treatment, cold hardening hours, cold acute stress exposure hours, and their interaction on larval-adult viability. Significance is defined as a p-value less than 0.05.

| **Response: Larval-adult viability** | **Chi-squre** | **Df** | **P-value (Chi-square)** |
|---|---|---|---|
| Selection treatment | 3.94998 | 2 | 0.139 |
| Hardening hours | 159.8652 | 2 | <0.001 |
| Acute stress exposure hours | 69.96492 | 2 | <0.001 |
| Selection treatment: hardening hours | 2.401897 | 4 | 0.662 |
| Selection treatment: acute stress exposure hours | 2.69116 | 4 | 0.611 |
| Hardening hours: acute stress exposure hours | 19.81748 | 4 | <0.001 |
| Selection treatment: hardening hours: acute stress exposure hours | 11.52897 | 8 | 0.173489 |

**Table S7**. Results of multiple comparisons on the effects of cold hardening hours and cold acute stress exposure hours on larval-adult viability. Significance is defined as a p-value less than 0.05.

| **Contrast Hardening hours at 0°C** | **Acute stress exposure hours at 0°C** | **Estimate** | **SE** | **Z ratio** | **P-value** |
|---|---|---|---|---|---|
| 0 - 1 | 4 | -0.239 | 0.082 | -2.913 | 0.004 |
| 0 - 2 | 4 | -0.357 | 0.072 | -4.973 | < 0.001 |
| 1 - 2 | 4 | -0.118 | 0.079 | -1.495 | 0.135 |
| 0 - 1 | 5 | -0.385 | 0.072 | -5.361 | < 0.001 |
| 0 - 2 | 5 | -0.485 | 0.081 | -6.020 | < 0.001 |
| 1 - 2 | 5 | -0.101 | 0.079 | -1.268 | 0.205 |
| 0 - 1 | 6 | -0.695 | 0.078 | -8.914 | < 0.001 |
| 0 - 2 | 6 | -0.723 | 0.078 | -9.331 | < 0.001 |

| | | | | | |
|---|---|---|---|---|---|
| 1 - 2 | 6 | -0.029 | 0.075 | -0.383 | 0.702 |
| **Acute stress exposure hours at 0°C** | **Hardening hours at 0°C** | **Estimate** | **SE** | **Z ratio** | **P-value** |
| 4 - 5 | 0 | 0.185 | 0.075 | 2.480 | 0.013 |
| 4 - 6 | 0 | 0.632 | 0.078 | 8.069 | <0.001 |
| 5 - 6 | 0 | 0.446 | 0.077 | 5.777 | <0.001 |
| 4 - 5 | 1 | 0.040 | 0.080 | 0.499 | 0.618 |
| 4 - 6 | 1 | 0.176 | 0.082 | 2.155 | 0.031 |
| 5 - 6 | 1 | 0.136 | 0.073 | 1.876 | 0.061 |
| 4 - 5 | 2 | 0.058 | 0.078 | 0.739 | 0.460 |
| 4 - 6 | 2 | 0.266 | 0.071 | 3.744 | <0.001 |
| 5 - 6 | 2 | 0.208 | 0.080 | 2.589 | 0.010 |

**Table S8**. Results of analysis of variance testing for effects of selection treatment, heat hardening hours, acute heat stress exposure hours, and their interaction on larval-adult viability. Significance is defined as a p-value less than 0.05.

| Response: Larval-adult viability | Chi-squre | Df | P-value (Chi-square) |
|---|---|---|---|
| Selection treatment | 4.484 | 1 | 0.034 |
| Hardening hours | 7.924 | 2 | 0.019 |
| Acute stress exposure hours | 76.147 | 2 | < 0.001 |
| Selection treatment: hardening hours | 719.448 | 2 | < 0.001 |
| Selection treatment: acute stress exposure hours | 11.881 | 4 | 0.018 |
| Hardening hours: acute stress exposure hours | 6.865 | 4 | 0.143 |
| Selection treatment: hardening hours: acute stress exposure hours | 110.150 | 4 | < 0.001 |

**Table S9**. Results of multiple comparisons on the effects of heat hardening hours, acute heat stress exposure hours on larval-adult viability. Significance is defined as a p-value less than 0.05.

| **Contrast** **Hardening hours at 35°C** | **Acute stress exposure hours at 39°C** | **Estimate** | **SE** | **Z ratio** | **P-value** |
|---|---|---|---|---|---|
| 0 - 1 | 0.25 | 0.106 | 0.092 | 1.156 | 0.248 |
| 0 - 2 | 0.25 | 0.401 | 0.091 | 4.380 | < 0.001 |
| 1 - 2 | 0.25 | 0.295 | 0.091 | 3.229 | 0.001 |
| 0 - 1 | 0.5 | -0.219 | 0.090 | -2.433 | 0.015 |
| 0 - 2 | 0.5 | -0.273 | 0.090 | -3.039 | 0.002 |
| 1 - 2 | 0.5 | -0.054 | 0.088 | -0.612 | 0.541 |
| 0 - 1 | 1 | -2.897 | 0.322 | -8.991 | < 0.001 |
| 0 - 2 | 1 | -2.919 | 0.323 | -9.040 | < 0.001 |
| 1 - 2 | 1 | -0.022 | 0.120 | -0.183 | 0.855 |

**Table S10**. Results of multiple comparisons on the effects of selection treatment, heat hardening hours, and acute heat stress exposure hours on larval-adult viability. Significance is defined as a p-value less than 0.05.

| **Selection Line = 18°C Standard diet** | **Estimate** | **SE** | **Z ratio** | **P-value** |
|---|---|---|---|---|
| **Acute stress exposure hours = 0.25** | | | | |
| Hardening hours = 0 -1 | 0.264 | 0.157 | 1.680 | 0.213 |
| Hardening hours = 0 -2 | 0.661 | 0.158 | 4.185 | <0.001 |
| Hardening hours = 1 -2 | 0.398 | 0.157 | 2.539 | 0.030 |
| **Selection Line = 18°C Standard diet** | | | | |
| **Acute stress exposure hours = 0.5** | | | | |
| Hardening hours = 0 -1 | -0.160 | 0.155 | -1.032 | 0.557 |
| Hardening hours = 0 -2 | -0.157 | 0.154 | -1.021 | 0.564 |
| Hardening hours = 1 -2 | 0.003 | 0.151 | 0.020 | 0.999 |
| **Selection Line = 18°C Standard diet** | | | | |
| **Acute stress exposure hours = 1** | | | | |
| Hardening hours = 0 -1 | -3.993 | 0.722 | -5.532 | <0.001 |
| Hardening hours = 0 -2 | -3.546 | 0.725 | -4.888 | <0.001 |
| Hardening hours = 1 -2 | 0.447 | 0.201 | 2.226 | 0.067 |
| **Selection Line = 25°C Standard diet** | | | | |
| **Acute stress exposure hours = 0.25** | | | | |
| Hardening hours = 0 -1 | 0.272 | 0.157 | 1.727 | 0.195 |
| Hardening hours = 0 -2 | 0.382 | 0.159 | 2.397 | 0.044 |
| Hardening hours = 1 -2 | 0.110 | 0.160 | 0.686 | 0.772 |
| **Selection Line = 25°C Standard diet** | | | | |
| **Acute stress exposure hours = 0.5** | | | | |
| Hardening hours = 0 -1 | -0.419 | 0.156 | -2.691 | 0.019 |
| Hardening hours = 0 -2 | -0.576 | 0.157 | -3.679 | <0.001 |
| Hardening hours = 1 -2 | -0.156 | 0.153 | -1.025 | 0.561 |
| **Selection Line = 25°C Standard diet** | | | | |
| **Acute stress exposure hours = 1** | | | | |
| Hardening hours = 0 -1 | -1.768 | 0.375 | -4.719 | <0.001 |
| Hardening hours = 0 -2 | -2.095 | 0.372 | -5.625 | <0.001 |
| Hardening hours = 1 -2 | -0.326 | 0.224 | -1.456 | 0.312 |
| **Selection Line = 28°C Standard diet** | | | | |
| **Acute stress exposure hours = 0.25** | | | | |
| Hardening hours = 0 -1 | -0.218 | 0.162 | -1.343 | 0.371 |
| Hardening hours = 0 -2 | 0.159 | 0.158 | 1.004 | 0.574 |
| Hardening hours = 1 -2 | 0.376 | 0.157 | 2.402 | 0.043 |
| **Selection Line = 28°C Standard diet** | | | | |

| | | | | |
|---|---|---|---|---|
| **Acute stress exposure hours = 0.5** | | | | |
| Hardening hours = 0 -1 | -0.078 | 0.157 | -0.497 | 0.873 |
| Hardening hours = 0 -2 | -0.086 | 0.156 | -0.551 | 0.846 |
| Hardening hours = 1 -2 | -0.007 | 0.151 | -0.051 | 0.999 |
| **Selection Line = 28°C Standard diet** | | | | |
| **Acute stress exposure hours = 1** | | | | |
| Hardening hours = 0 -1 | -2.919 | 0.523 | -5.607 | <0.001 |
| Hardening hours = 0 -2 | -3.116 | 0.523 | -5.957 | <0.001 |
| Hardening hours = 1 -2 | -0.187 | 0.199 | -0.940 | 0.615 |

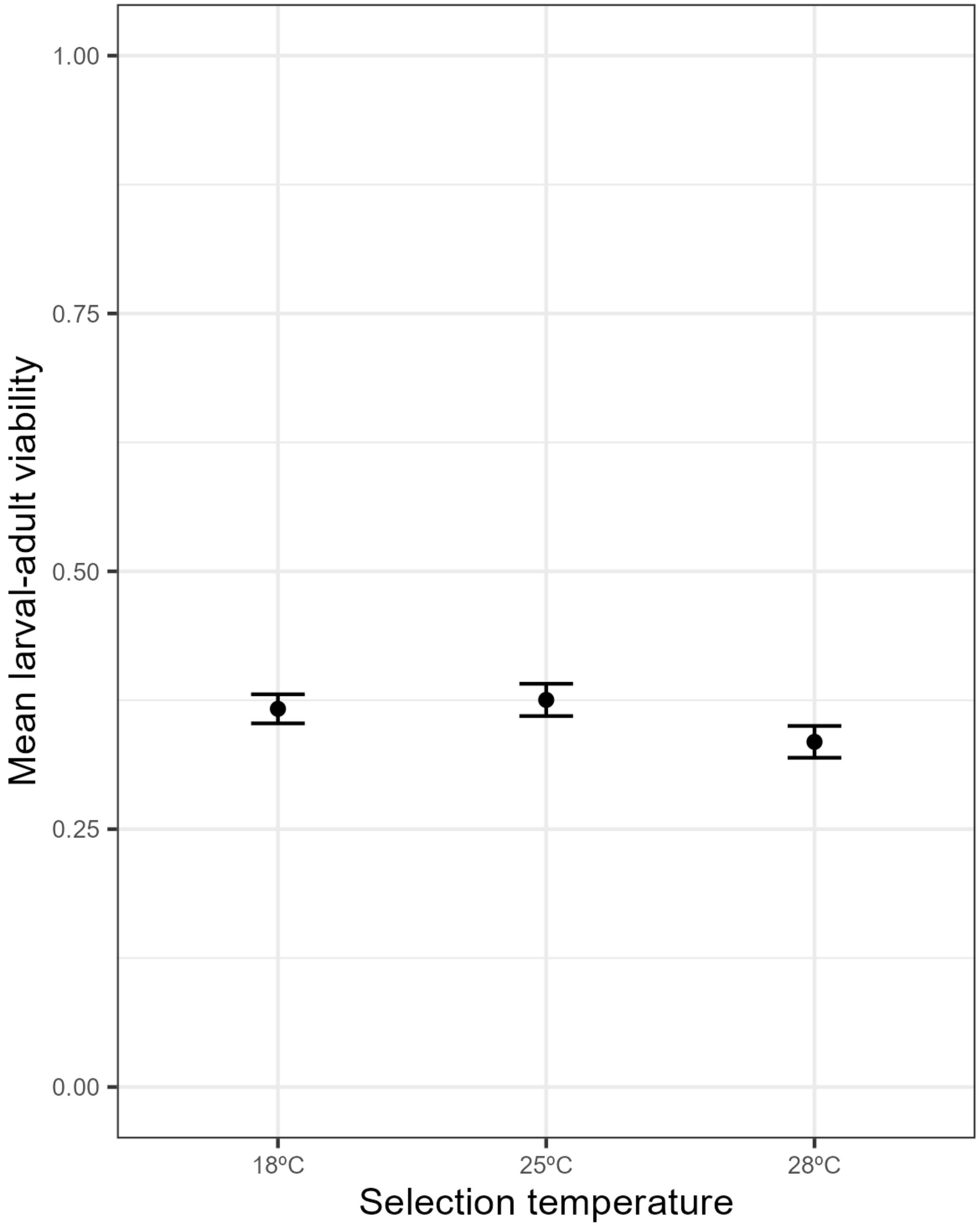


**Figure S1**. Mean larval-adult viability of thermal selection lines after exposure to an acute heat shock treatment of 0.5 h at 39°C, averaged across selection diet, hardened and non-hardened larvae. Error bars indicate ± 1 standard error.

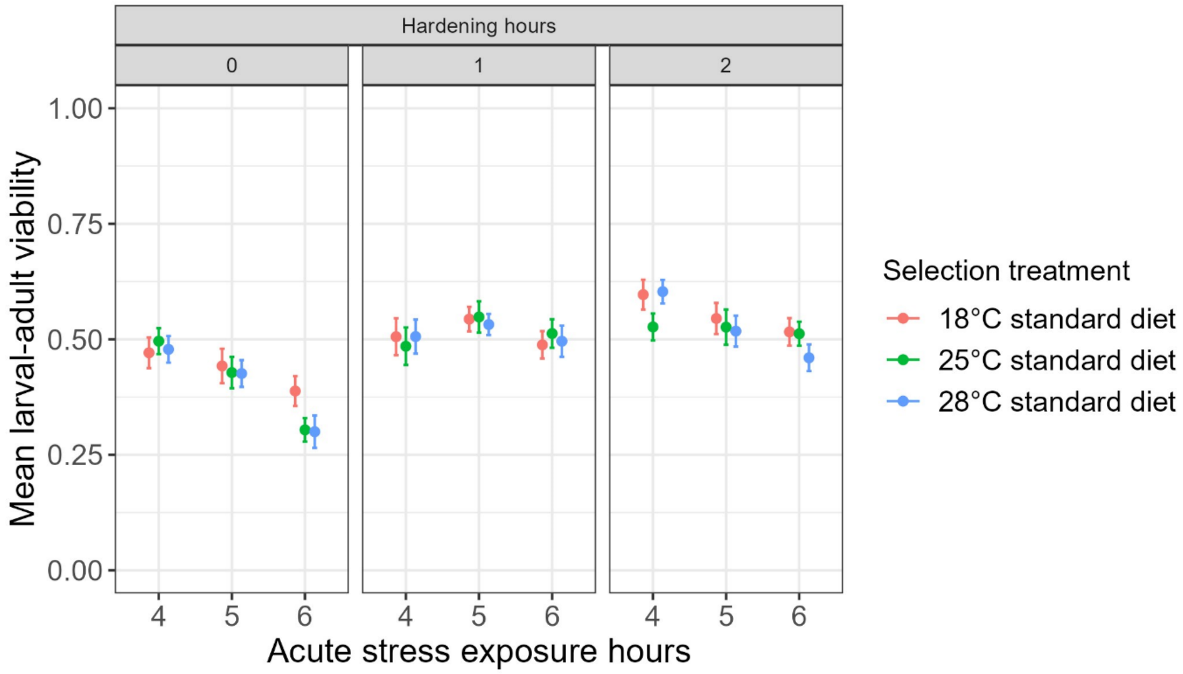


**Figure S2**. The effects of different durations of 0°C cold hardening treatment and 0°C acute cold stress treatment on mean larval-adult viability of experimental evolution lines. Error bars indicate ± 1 standard error.

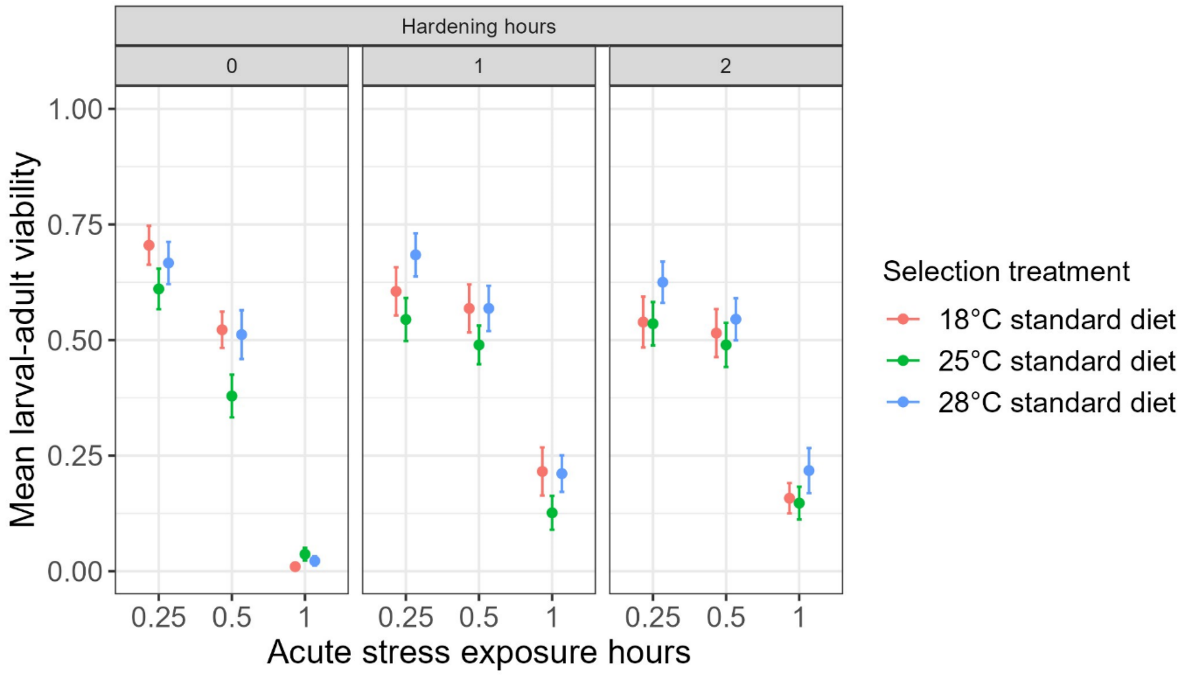

**Figure S3**. The effects of different durations of 35°C heat hardening treatment and 39°C acute heat stress treatment on mean larval-adult viability of experimental evolution lines. Error bars indicate ± 1 standard error.
